# Stochastic analysis of optimal production of infectious progeny in Chlamydia

**DOI:** 10.1101/836353

**Authors:** German Enciso, Ming Tan, Frederic Y.M. Wan

## Abstract

Recent data collected on the Chlamydia Trachomatis life cycle show an initial period of no RB-to-EB conversion. This and other features of the observed bacterial life cycle are postulated to be consequences of the bacteria’s drive for Darwinian survival. Stochastic optimal control models formulated herein in fact lead to an initial conversion holiday that support this proposition.

## 1. THE LIFE CYCLE OF C. TRACHOMATIS

Chlamydia trachomatis is a bacterium that causes ocular and genital tract infections in humans. In ocular infections, *C. trachomatis* is the world’s leading cause of preventable blindness. This condition, known as trachoma, currently affects 84 - 150 million people in the world, causing blindness to 8 million people. It has been targeted for elimination by the World Health Organization. (For references on data related to this infectious disease, see [5, 3] and references cited therein)

As a pathogenic bacterium, *C. trachomatis* has an unusual intracellular developmental cycle involving conversion between Chlamydial forms within a cytoplasmic inclusion. The simplest model has the bacteria in two forms [3]: a *reticulate body* (RB) form of the bacteria that repeatedly divides by binary fission and asynchronously differentiates into the infectious *elementary body* (EB) form that does not divide. Upon the lysing of the host cell, only the EB form of the bacteria survives and proceeds to infect other cells. Once an EB enters a new host cell, it turns into an RB and begins to replicate, repeating the life cycle as shown in Figure 1. We abbreviate the event of the initial EB form of the bacteria entering the host cell and immediately converting into RB form as the “*initial infecting RB population*” and denote it by *N* (the number of initial RB units). The model allows us to examine the optimal conversion of some or all RB units into EB at each instant of time *t* to maximize the EB population at a *terminal time T* when the host cell lyses. The optimization in our optimal control model constitutes an effort to uncover the possibility of a Darwinian force that drives the life cycle of the bacteria.

**Figure 1:** Life cycle of Chlamydia Trachomatis.

Until recently, these developmental events of Chlamydia life cycle have not been quantified because conventional electron-microscopy only visualizes sections of the large Chlamydial inclusion. Using a novel three-dimensional electron microscopy (EM) approach (known as serial block-face scanning EM (SBEM) technique), the lab of Tan and Suetterlin obtained recently extensive quantitative data on RB replication and RB-to-EB conversion during the developmental cycle [3]. Among the principal findings is the existence of an initial period of no RB-to-EB conversion after the initial infection of a healthy host cell. It is our conjecture that such an unusual life cycle is the consequence of the usual Darwinian force at work to maximize the spread of the infectious disease. In our first effort test this proposition, we worked with deterministic optimal control models for a prescribed (known) terminal time for the life cycle and investigated consequences of several proliferation and conversion scenarios [5]. However, empirical data in [3] show variations of life cycle features such as terminal time and initiation of the RB-to-EB conversion. These findings prompted us to initiate a few stochastic models to address these variations.

We begin with a simple birth and death process model for the evolution of the RB and EB populations that focuses on the complementary choice of division of RB (birth) and conversion of RB to EB (death) over a finite time *T* before the host cell lyses. A more realistic feature (than a prescribed terminal time for the models in [5]) toward the same goal of determining the maximal terminal infectious EB population is a criterion for setting the terminal time of the proliferation-conversion cycle based on the empirical findings of [3]. In the first model, we consider only time invariant probabilities for division and RB-to-EB conversion. An optimal division-to-conversion probability ratio is found for maximizing the expected EB population at terminal time subject to a threshold of weighted sum of the (expected) RB and EB populations at the terminal time.

The optimal conversion strategy of this first model can clearly be improved by a more general model that allows for a time-varying conversion probability with the expectation of a larger expected terminal EB population than that induced by a time-invariant conversion-to-duplication probability ratio. The optimal conversion strategy of this second model shows that maximum terminal EB population is in fact attained by a bang-bang conversion strategy. This result of the models in [5] is remarkable not only for its agreement with the empirical finding of a *period of no conversion* at the start of a new life cycle reported in [3], but also for buttressing the proposition that the *conversion holiday* is a consequence of the Darwinian force at work for the optimal proliferation and survival of the bacteria.

This second model still needs further modification to accurately reflect the continual growth of the RB population at least for a period after the initiation of RB-to-EB conversion shown in the data reported in [3]. A further refined model that allows for multiple forms of both RB and EB as observed in [3] is then shown to have a period of RB growth after the start of conversion..

To address the observed variation of terminal time of the life cycle, we formulate and analyze a fourth probabilistic model where the only knowledge of the uncertain terminal time is a prescribed *probability density function* (pdf). The maximization of the expected terminal EB population results in a typical stochastic optimization problem. It leads again to a bang-bang conversion strategy, showing that an uncertain terminal time does not affect the proposition that a Darwinian force is at work in shaping the life cycle of the bacteria.

## 2. A BIRTH AND DEATH PROCESS MODEL

### 2.1. The Kolmogorov Equations

To model RB proliferation (birth) and RB-to-EB conversion (death) of the Chlamydia life cycle as a birth and death process, we denote by *R*(*t*) and *E*(*t*) the random variable for the size of *RB* and *EB* population, respectively, at time *t*. Let *P*_*k*_(*t*) and *Q*_*j*_(*t*) be the probability of *R*(*t*) and *E*(*t*) be of size *k* and *j*, respectively, at time *t*. With assumptions similar to those for various birth and death models (such as stationarity, independent increments and a sufficiently short elapsed time *δt* for no more than one division or conversion), we have the following relations for *P*_*k*_(*t* + *δt*) and *Q*_*j*_(*t* + *δt*):

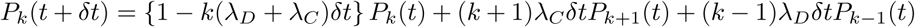

and

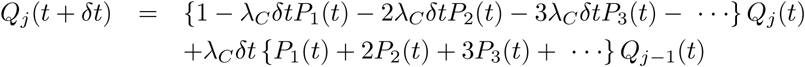

except for negligibly small terms relative to terms proportional to *δt*. In these two relations, *λ*_*C*_ *δt* and *λ*_*D*_*δt* are the probability of one conversion and one duplication, respectively, during the elapsed time *δt*. As a first model (that is to be refined later), we let both *λ*_*C*_ and *λ*_*D*_ be independent of time and population size.

In the limit as *δt* tends to zero, we get

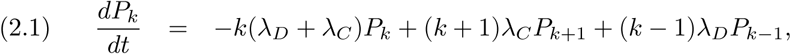

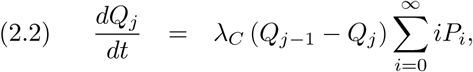

augmented by the initial conditions *P*_*k*_(0) = *δ*_*kN*_ and *Q*_*j*_(0) = 0, *k, j* = 0, 1, 2, …, given that the process starts with *N* RB units and no EB.

### 2.2. Solution by Generating Functions

The ODE system for *P*_*k*_(*t*) can be reformulated into a problem for the corresponding generating function Σ*P*_*k*_(*t*)*x*^*k*^ = *G*(*x, τ*)

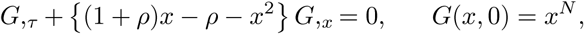

with

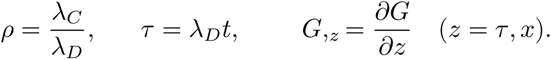

The IVP for this first order PDE is solved by the method of characteristics to give for *ρ* ≠ 1

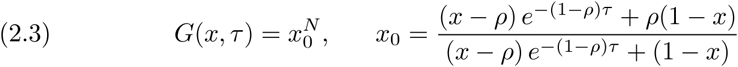

Though it can be obtained as a limit of (2.3), the special case of *ρ* = 1 (*λ*_*C*_ = *λ*_*D*_) will be derived separately in the next subsection.

Note that

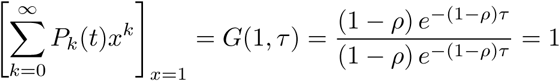

and

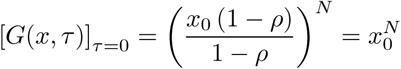

as they should be. More importantly, we have the following expression for the *expected value* of the random variable *R* (*t*) ≡ *R*_*t*_ for a fixed *t* needed for the determination of the generation function for the {*Q*_*j*_(*t*)}:

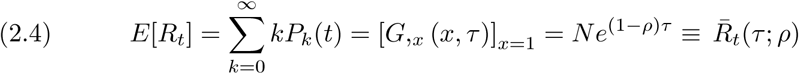

with

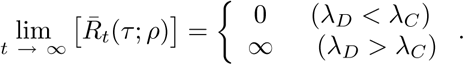

The expressions *E*[*R*_*t*_] and 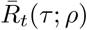 will be used interchangeably for the expected RB population size as a function of time *t* with *τ* = *λ*_*D*_*t* and *ρ* = *λ*_*C*_ /*λ*_*D*_. Similar expressions will also be used for the expected EB population size below.

Turning now to the ODE system for {*Q*_*j*_(*t*)}, we also formulate the IVP for the corresponding generating function *F* (*x, τ*) = Σ *Q*_*j*_(*t*)*x*^*j*^ to get

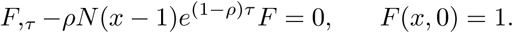

The equation for *F*(*x, t*) above is a first order linear ODE that can be solved with the help of an integrating factor to get

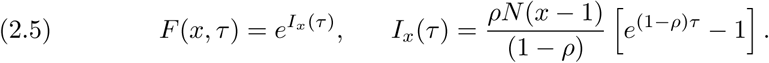

### 2.3. The Special Case λ_C_ = λ_D_

For *ρ* = 1, the IVP for the generating function *G*(*x, τ*) becomes

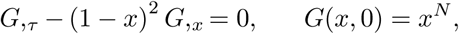

with

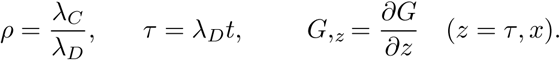

Its solution by the method of characteristics is

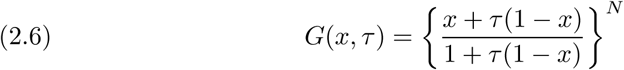

with

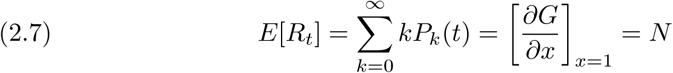

needed for the generating function of {*Q*_*j*_(*t*)}. Note that *E*[*R*_*t*_] for *ρ* = 1 is time-invariant.

The IVP for *F* (*x, τ*) = Σ*Q*_*j*_(*t*)*x*^*j*^ for *ρ* = 1 is

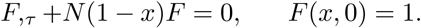

Its solution is

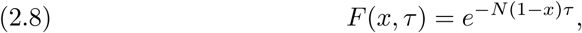

which can also be deduced as the limit of (2.5) as *ρ* → 1.

### 2.4. Means, Variances and Distributions

#### 2.4.1. Distributions

We can obtain {*P*_*k*_(*t*)} and {*Q*_*j*_(*t*)} by expanding *G*(*x, τ*) and *F*(*x, τ*), respectively, as Taylor series in *x* with

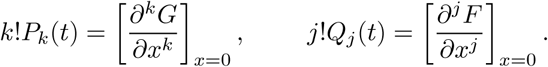

The simpler distributions for {*Q*_*k*_(*t*)} of primary interest are easily calculated from (2.5). For *ρ* ≠ 1, we have

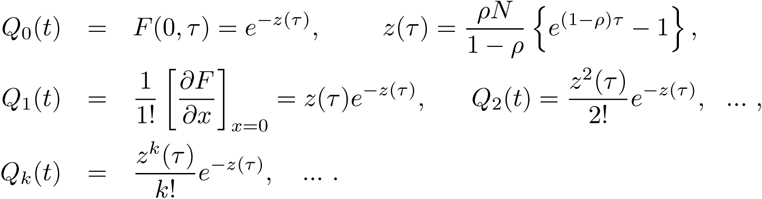

Evidently, the distributions for the random variable *E*_*t*_ is Poisson in the composite time-dependent variable *z*(*τ*). This is further supported by the statistics for *E*_*t*_ found below where its variance is identical to its mean.

From the more complicated generating function *G*(*x, τ*), we obtain

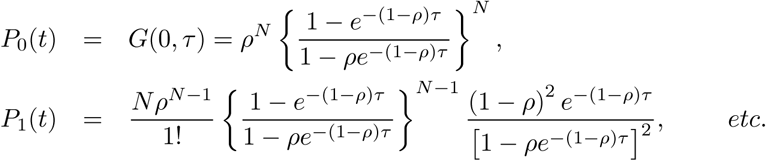

For *ρ* > 1 (so that conversion is at a faster rate than the natural growth rate of the RB population), we have the following limiting behavior for the available expressions for *P*_0_(*t*) and *P*_1_(*t*)

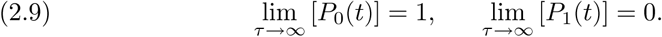

The expected steady state behavior of the remaining probability distributions {*P*_*k*_(*t*)}, *k* > 1, can be obtained directly from the corresponding steady state Kolmogorov equations

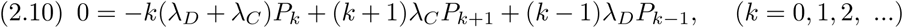

For *k* = 0 and *k* = 1, the master equation above reproduces the results in (2.9). These in turn require

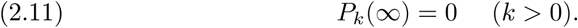

This steady state behavior is consistent with the eventual complete conversion of RB to EB if the conversion is allowed to continue indefinitely.

For *ρ* = 1, it is known from previous calculations *E*[*R*_*t*_] = *N*. This would certainly require *P*_*k*_(*t*) → 0 as *k* → ∞. The exact solution (2.6) in fact requires from or

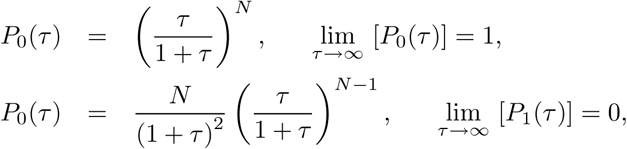

etc. In fact, if a limiting steady state solution exists for *ρ* = 1, the steady state Kolmogorov equation (2.10) also requires (2.11) to apply with *P*_0_(∞) = 1.

For *ρ* < 1, the distributions do not reach a steady state since RB populations continues to grow indefinitely. Altogether, the distributions for *R*_*t*_, transient or steady state (if it exists) are seen not associated with the Poisson distribution.

#### 2.4.2. Means

Of more immediate interest is the expected values for the random variables *R*_*t*_ and *E*_*t*_ for the RB and EB population, respectively, at a fixed *t*, denoted by 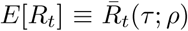 and, *E*[*E*_*t*_] ≡ *Ē*_*t*_(*τ*; *ρ*) respectively. These are denoted by We have already found the mean of *R*_*t*_ in (2.4) to be

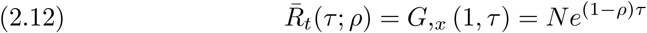

which holds for *ρ* = 1 as well. The corresponding expected value *E*[*E*_*t*_] = *Ē*_*t*_(*τ*; *ρ*) = *F*,_*x*_ (1,*τ*) of the EB population is found from (2.5) and (2.8) to be

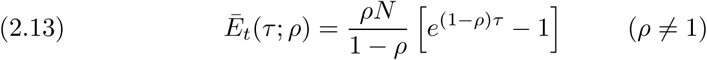

and

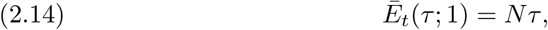

respectively. The latter can also be obtained as a limiting case of (2.13) as *ρ* → 1.

#### 2.4.3. Variances

For the corresponding variances, we note

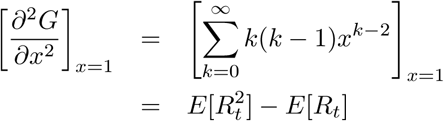

so that

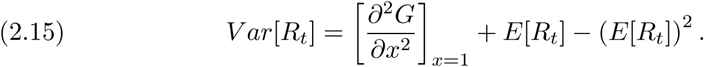

For *ρ* ≠ 1, we have from (2.3) and (2.12)

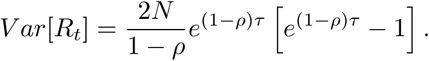

The limiting case of *λ*_*C*_ = *λ*_*D*_ can be obtained by taking the limit of the above expression as *ρ* → 1 to get

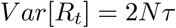

(and confirmed independently by using the results (2.6) and (2.7) in (2.15)). In either case, the variance is not equal to the mean with the latter independent of time.

By similar calculations, we get from (2.5) and (2.8) the corresponding variance

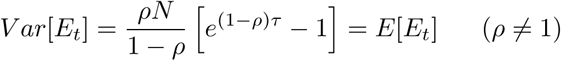

with the limiting value *Nτ* as *ρ* → 1. Unlike the statistics of the RB population, the variance of EB population is equal to the mean which is as expected given the distributions for *E*_*t*_ already obtained above.

## 3. MAXIMUM EXPECTED EB POPULATION AT HOST CELL LYSIS TIME

### 3.1. Terminal Time of Life Cycle

Data collected in [3] shows that the total chlamydia units in an inclusion typically asymptote to an upper limit after 32 hpi. The inclusion volume also increases with time at the early stage of the developmental cycle but also ceases to increase near the end. It was found that the total volume (size) of the chlamydiae actually reduces while the total number of units increases, available space for more Chlamydia does not appear to be an issue. Possible cause for host cell lysing to end the life cycle will be discussed later in this section. The empirical data would suggest the following relation among the *expected* RB and EB population at the terminal time *T* of the developmental cycle:

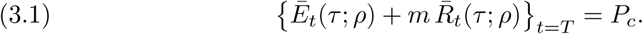

where *P*_*c*_ is the total chlamydiae asymptote. For the purpose of analysis, we consider the terminal time of a particular life cycle; the variation of *T* among the different host cells will be treated in the last two section of this paper. For the case *m* > 1 reflecting the relative size of the two types of bodies (with *m* as high as 50 immediately after infection but decreasing to about 5 near the end of the life cycle). The analysis for this specific choices of *m* (suitably modified) will be shown to apply to other choices of *m* (including 0 < *m* < 1 and *m* < 0) that are both meaningful and realistic.

In the subsequent development, we take the Chlamydia life cycle to be the consequence of the bacteria’s strive for Darwinian survival and the conversion probability is evolved from the bacteria’s continuing effort to maximize the terminal expected EB population. In that context, the parameters *λ*_*D*_ and *P*_*c*_ are known (or estimated from available data) and *N* is prescribed (taken to be 1 in control experiments). The RB-to-EB conversion probability *λ*_*C*_ is taken to be the instrument for maximizing the terminal EB population for optimal spread of the bacteria. The relation (3.1) may then be taken as a condition for determining the terminal time *T* as a function of the conversion-to-duplication probability ratio *ρ* = *λ*_*C*_ /*λ*_*D*_. Even for this one free parameter case, it is still rather tedious to try to solve for *T* in terms of *ρ* (with *N* and *λ*_*D*_ fixed). In determining the maximum point *ρ*_max_ that maximizes the expected EB population at terminal time *T*, our approach is to determine *ρ*_max_ and the corresponding normalized terminal time *τ* _max_ = *τ*_*T*_ (*ρ*_max_) simultaneously.

### 3.2. Maximum Point for Expected Terminal EB Population

With *N* and *λ*_*D*_ prescribed and *τ*_*T*_ = *λ*_*D*_*T*, the problem of maximizing *Ē*_*t*_(*τ*_*T*_; *ρ*) subject to the constraint (3.1) will be solved by the method of Lagrange multiplier. By this approach, we seek a maximum point 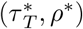 of the augmented objective function

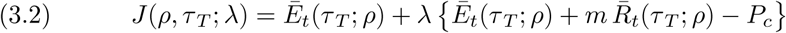

(subject to the constraint (3.1)) where *λ* is a Lagrange multipliers.

Any maximum point must be a stationary point of (3.2) with

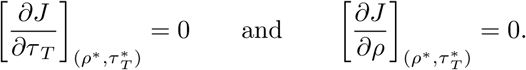

The first condition requires

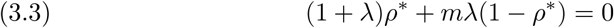

since 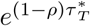 does not vanish. Upon observing (3.3), the second condition requires

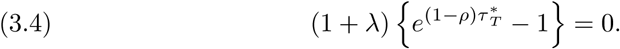

since 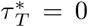 is not acceptable (being a minimum point with *Ē*_*t*_ = 0), the two conditions (3.3) and (3.4) yield a single stationary point

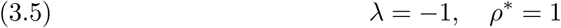

with 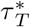 determined by (3.1) to be

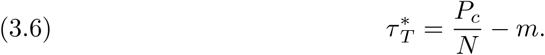

### 3.3. The Maximum Expected EB Population

The corresponding maximum expected EB at terminal time is then

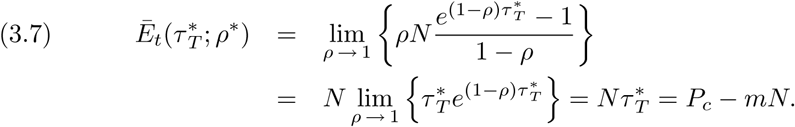

We highlight this outcome in the following proposition:

#### Proposition 1.

*For a prescribed combination of λ*_*D*_ *and N, the optimal (time-invariant) RB-to-EB conversion probability is* 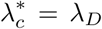 *for a maximum expected terminal EB population of P*_*c*_ − *mN attained at the terminal time*

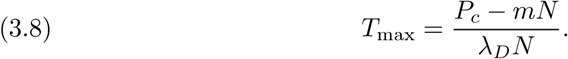

#### Remark 1.

*Before leaving the completed solution for the present birth and death process model, the following two observations are appropriate at this time:*

- *The requirement of optimal conversion probability* 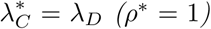 *may not be met if the conversion capacity is limited so that λ*_*C*_ *is restricted to be* 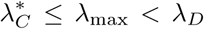. *If λ*_max_ < *λ*_*D*_, *we should presumably convert (suboptimally) at* 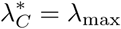.
- *The rather restrictive model with of a time invariant conversion probability λ*_*C*_ *serves only a stepping stone to a more appropriate model with a time-varying alternative since the larger set of admissible comparison functions would generally lead to a larger (expected) terminal EB population*.

It turns out that a simple (but qualitatively significant) change to a time dependent *λ*_*C*_ (*t*) not only leads to an optimal conversion rate that agrees qualitatively with available data over a substantial period after infection but also a faster host cell lysis time that is comparable to the observed terminal time of the life cycle.

### 3.4. On the Terminal Time Condition

Before we embarked on seeking a more appropriate model than one with time-invariant conversion, we should discuss further the terminal time condition (3.1) as indicated previously. That condition for determining the terminal time is a special case of the more general condition

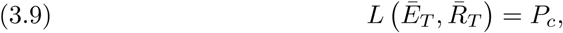

where

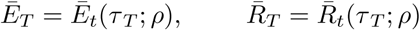

and *L* (·, ·) is some differentiable function of two variables. As long as

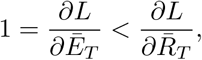

there would generally be no significant qualitative change in the results due to the more general condition (3.9). Here, we examine instead other possible modifications of the threshold condition that are qualitatively different from (3.1).

The condition (3.1) with *m* > 1 may be associated with the relative size of the EB and RB particles (by a factor of 50 at the start and down to about 5 near the end). Experimental findings reported in [3] also suggest that there are additional factors contributing to the lysing of the host cell. Among these is EB particles secreting chemicals that enhance host cell lysing. To incorporate this additional effect, we may again limit the threshold criterion to a linear condition of the form (3.1) for a fixed total Chlamydia population threshold *P*_*c*_ but modify it to read

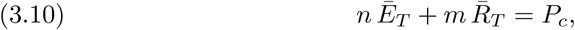

with the factor *n* > 1 characterized the effects of the secreted chemicals. Depending on the potency of the lysis inducing EB chemicals, the ratio *m*/*n* > 0 may be > 1 or < 1. In that case, the results obtained for (3.1) again applies but with *m* and *P*_*c*_ replaced by *m*/*n* and *P*_*c*_/*n*. In particular, we have as the maximum expected EB,

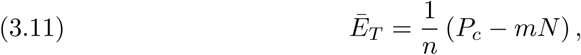

to be attained at the terminal time

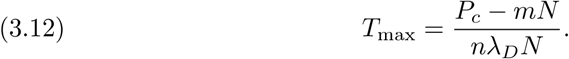

With *n* > 1, we see from (3.8) and (3.11) that the terminal time would be sooner with a smaller expected terminal EB population.

Observations in [3] also suggest the possibility of RB particles exhibiting inhibitory effects on host cell lysing to prolong their own existence. In the context of the linear threshold condition (3.10), such inhibitory effects may be incorporated by modifying (3.10) to read

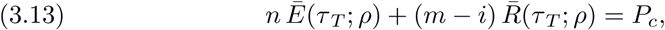

for some *i* > 0. The results (3.11) and (3.12) again apply with *m* replaced by *m* − *i*. It follows that for a given pair of (*m, n*) values, the terminal time *T* would be later and the expected terminal EB population would be larger.

When warranted, the incorporation of the two new effects into the lysing threshold condition (3.1) can again be extended to the nonlinear form (3.9) with suitable conditions on the partial derivatives of *L* with respect to *E*_*T*_ and *R*_*T*_, respectively. Our more immediate task however is to seek a more superior conversion strategy to result in a larger expected terminal EB population by allowing *λ*_*C*_ (or equivalently *ρ*) to vary with time.

## 4. BIRTH AND DEATH MODEL WITH TIME-VARYING CONTROL

### 4.1. Time-dependent Conversion Probability

The simple birth and death type probabilistic model of the last section provides a stepping stone to a more appropriate models of the same type. For such second model, we only need to make a simple change from the first model by asking whether or not we may attain a faster spread of the infectious bacteria by allowing the conversion probability to vary with time. To initiate an effort in this direction, we continue to work with (2.1) and (2.2) but allow *λ*_*C*_ to depend on *t*. Upon re-writing them in term of normalized time *τ* = *λ*_*D*_*t*, we have the following Kolmogorov ODE for the new model:

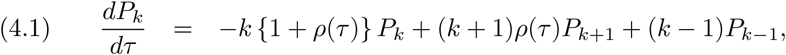

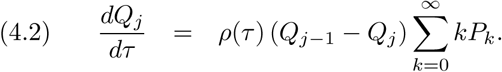

Similar to the time-invariant case, they give rise to the PDE

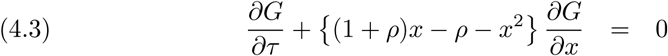

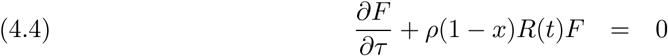

for the two pgf

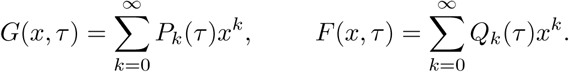

The initial conditions for {*P*_*k*_(*τ*)} and {*Q*_*j*_(*τ*)} lead again to

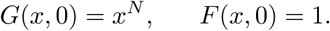

### 4.2. Application of the Method of Characteristics

The characteristic ODE for the first order PDE for *G*(*x, τ*) are

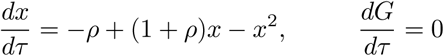

The ODE for *x*(*τ*) is a Riccati equation for which there is no general method of solution when *ρ* varies with time. For our particular right hand side, it is possible to factor the quadratic expression to give

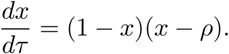

Evidently, *x*_*p*_(*τ*) = 1 is a solution of the ODE. It is only a *particular* solution since it does not contain any constant of integration to fit a prescribed auxiliary condition. However, if we let

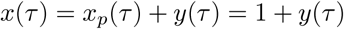

and substitute it into the original Riccati equation, we obtain

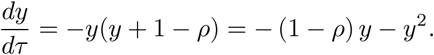

The first order ODE for *y*(*t*) is a Bernoulli equation. It is transformed by *y* = 1/*z* into the linear ODE

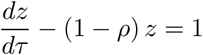

with the exact solution

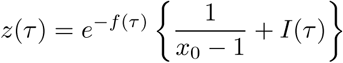

where

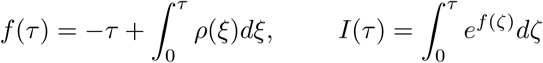

The corresponding solution for *x*(*τ*) is

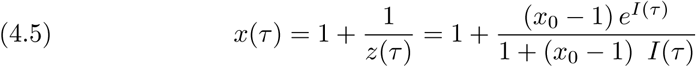

with

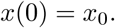

The solution for the other characteristic ODE is.

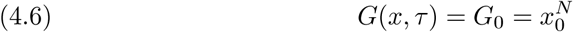

since there are exactly *N* RB units initially so that

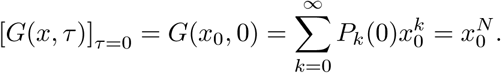

To complete the solution, we solve (4.5) for *x*_0_ in terms of *x* and *τ* to get

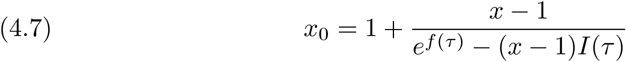

Upon using this expression for *x*_0_ in (4.6), we obtain

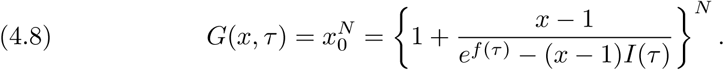

The expected RB population as a function of time is then given by

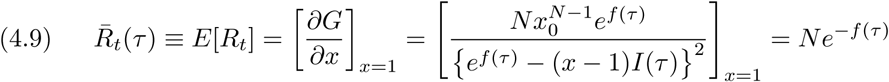

with [*x*_0_]_*x*=1_ = 1 (see (4.7)).

For the special case of a time-invariant *ρ*, the solution (4.8) and (4.9) reduce to (2.3) and (2.4), respectively.

### 4.3. The Expected EB Population

Unlike the PDE (4.3) for the generating function *G*(*x, τ*) of the sequence {*P*_*k*_(*τ*)}, the first order ODE (4.4) for the generating function *F* (*x, τ*) of the sequence {*Q*_*k*_(*τ*)} is separable in time with *x* as a parameter. With 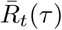 given by (4.9), (4.4) becomes

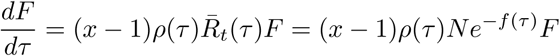

so that

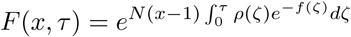

with *F* (*x*, 0) = 1 meeting the condition of no EB initially.

The expected EB population at any instant in time is then

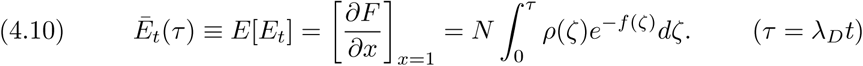

For a constant *ρ*, the expression for *E*(*t*) simplifies to the result obtained earlier in (2.13) for this special case.

## 5. MAXIMUM TERMINAL EB POPULATION WITH TIME-VARYING CONVERSION

### 5.1. Optimal Control and the Maximum Principle

As indicated earlier, we wish to find the *ρ*(*τ*) that maximizes the expected terminal EB population *Ē*_*T*_ ≡ *Ē*_*t*_(*τ*_*T*_) with *τ*_*T*_ = *λ*_*D*_*T*. It is well-known that the appropriate method of solution for this type of problems is the method of optimal control. To apply this method, we henceforth change to the more conventional optimal control notations, writing *u*(*t*), *α, E*_*T*_, *E*(*t*) and *R*(*t*) for *λ*_*c*_(*t*), *λ*_*D*_, *Ē*_*T*_, *Ē*_*t*_(*τ*) and 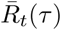, respectively, as they allow for direct comparison with the results for the deterministic optimal control problem of [5, 7]. In terms of the new notations, we revert to the original un-normalized quantities and state the optimization problem as

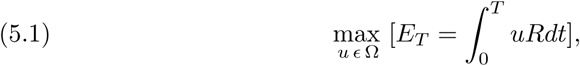

with the optimal control *u*_*op*_(*t*) to be chosen among all *u*(*t*) (≡ *λ*_*C*_ (*t*)) in the admissible set Ω of piecewise smooth (PWS) functions. The maximization is subject to the growth dynamics and initial condition

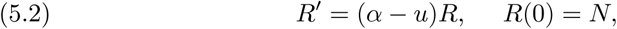

(equivalent to the exact solution (4.9) and (4.10)) and the terminal constraint

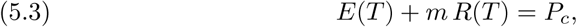

with *R*_*T*_ = *R*(*T*) is the (expected) terminal RB population in the new notation. (To be concrete, we will limit our discussion to the case *m* > 1. it will be obvious when the results obtained by our analysis may apply (beyond the basic relative size consideration) to the more general case with EB promoting lysis while RB inhibiting lysis.)

Given the reality of a limited rate for the one-way RB-to-EB conversion, the optimal choice *u*_*op*_(*t*) *ϵ* Ω is required to satisfy the inequality constraints

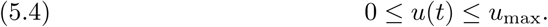

The following proposition relates the growth dynamics (5.2) to the probabilistic model of the previous chapter and establishes the consistency of the two different formulations for determining the expected RB and EB populations as functions of time.

#### Proposition 2.

*The rates of growth of the expected RB population R*(*t*) *and the expected EB population E*(*t*) *are determined by the IVP (5.2) and*

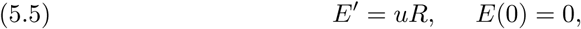

*respectively*.

*Proof*. Upon multiplying (2.1) by *k* and summing from 0 to ∞, we obtain

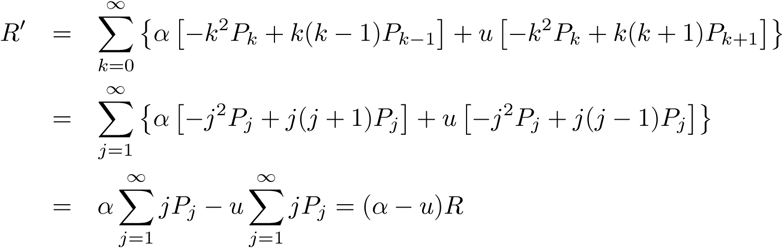

proving the validity of the ODE in (5.2). The initial condition in (5.2) follows from

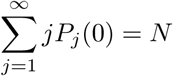

given *P*_*j*_(0) = *δ*_*jN*_. The second IVP can be derived similarly from (2.2) and *Q*_*j*_(0) = *δ*_*j*0_. □

### 5.2. The Maximum Principle

Our goal is find the optimal conversion rate *u*_*op*_(*t*) that maximizes the expected terminal EB population *E*_*T*_ by application of the Maximum Principle [1, 6, 4]. For this method of solution, we introduce the Hamiltonian

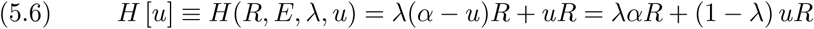

In terms of the Hamiltonian, the Maximum principle requires the following adjoint differential equation

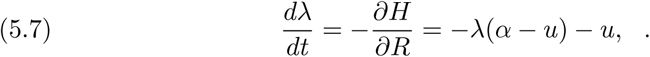

be satisfied by the adjoint function (aka Lagrange multiplier) *λ*(*t*) associated with the state equation in (5.2) for *R*(*t*). A condition on the unknown *R*(*t*) is prescribed only at *t* = 0 of the solution domain [0, *T*]. (While the threshold condition (5.3) related *R*(*T*) to *E*(*T*), the latter is to be maximized and hence also not prescribed.) The Maximum Principle then requires the adjoint function *λ*(*t*) to satisfied the adjoint (Euler) boundary condition

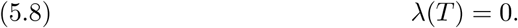

If the control *u*(*t*) should have a finite jump discontinuity at an instant *t*_*s*_ in (0, *T*), the Maximum Principle requires that the Hamiltonian be continuous at the *switch point t*_*s*_:

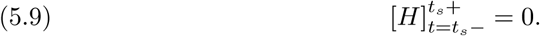

Finally, the *optimal control u*_*op*_(*t*) must maximize *H*(*t*) in the sense of

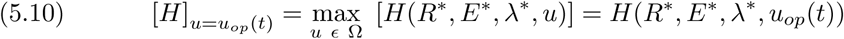

where ()* is the quantity () induced by the optimal control. As we shall see, this last step constitutes the most challenging part of the solution process for our problem.

The second order ODE system (5.2) and (5.7) is supplemented by two auxiliary conditions: the initial condition in (5.2) and the Euler boundary condition (5.8). Together, they determine the state function *R*(*t*) and adjoint function λ(*t*), with the terminal time *T* determined by the threshold condition (5.3), once we know the control *u*(*t*). If the control should have a finite jump discontinuity at a switch point *t*_*s*_, the switch condition (5.9) applies to determine the switch point.

It remains apply (5.10) to specify the optimal control *u*_*op*_(*t*) to complete the solution process. From the requirements imposed by the Maximum Principle, we readily observe in the following three sections three key results pertaining to the optimal conversion strategy, the *optimal control u*_*op*_(*t*).

### 5.3. Singular Solution Not Applicable

Suppose we have the state and adjoint functions corresponding to the optimal control *u*_*op*_(*t*). Then the dependence of the Hamiltonian on the control must be such that *u*_*op*_(*t*) is one of its maximum points. We should therefore seek *u*_*op*_(*t*) among the stationary points *u*_*S*_(*t*) of the Hamiltonian *H* [*u*(*t*)]. By the condition (5.10) of the Maximum Principle, this stationary condition requires

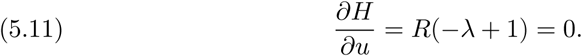

The relation (5.11) does not involve the control *u*(*t*) and therefore does not provide any clue to *u*_*S*_(*t*) directly. With *R*(*t*) > 0, the stationary condition (5.11) can be met by the *singular solution*

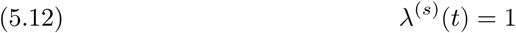

so that the corresponding adjoint function *λ*^(*s*)^(*t*) is independent of time.

In principle, the singular solution (5.12) for the adjoint functions should be preferred since it renders the Hamiltonian stationary and may correspond to a maximum point. However, it is not appropriate for any sub-interval of the solution domain. If the singular solution should be optimal in some subinterval (*t*_1_, *t*_2_) of [0, *T*], then the adjoint DE for *λ* (5.7) requires

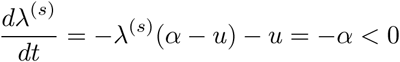

for any time *t* in (*t*_1_, *t*_2_). This contradicts the fact that [*λ*^(*s*)^]′ = [1]′ = 0; hence, *λ*^(*s*)^(*t*) is not optimal in (*t*_1_, *t*_2_). Since the observation holds for any subinterval of [0,*T*], we have established the following negative result for our problem:

#### Proposition 3.

*The singular solution (5.12) does not apply to (and therefore has no role in) our model of maximum terminal EB population with a time-varying control*.

### 5.4. Maximum Conversion for *u*_max_ ≤ *α*

#### Proposition 4.

*If u*_max_ ≤ *α (henceforth written as u*_*α*_ ≡ *u*_max_ − *α* ≤ 0*), we must have*

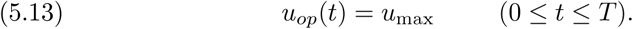

*Proof*. If *u*(*t*) < *u*_max_, we can increase *E*(*T*) by converting at *u*_max_ at any time *t* and still continue to grow the RB population. □

The corresponding exact solutions for the RB and EB populations for *u*_*op*_(*t*) = *u*_max_ are

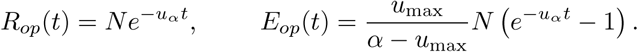

The terminal time *T* is determined by the constraint (5.3) to be.

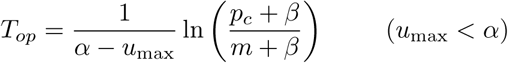

where

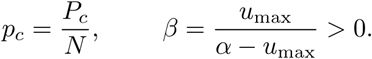

With *p*_*c*_ ≫ *m* (and *u*_max_ < *α*), *T*_*op*_ > 0 is assured. The corresponding terminal EB population is

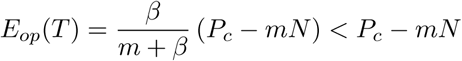

as we would expect (since *u*_*op*_(*t*) = *α* is known to be optimal from the time-invariant control analysis).

As *u*_max_ ↑ *α*, we have

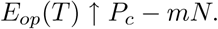

After application of L’Hospital’s rule, the companion limiting value for the terminal time is found to be

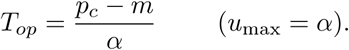

Both are the same as those found for a time-invariant *u* = *λ*_*C*_ earlier.

In all cases, the results apply for *m* < *P*_*c*_/*N* = *p*_*c*_. For the less likely case of *m* > *P*_*c*_/*N*, the requirement (5.3) is already met at (or prior to) the initial time *t* = 0 without any EB units. Either the host cell cannot violate the terminal time condition or the condition *u*_max_ ≤ *α* does not apply. The latter will be further supported by the principal development below for the present model with time-varying conversion rate.

### 5.5. Bang-Bang Control for *α* < *u*_max_

For the optimal control in this range of *u*_max_, we first observe the following properties for the solution of the problem:

#### Property 1

For *u*_*α*_ > 0, we must have

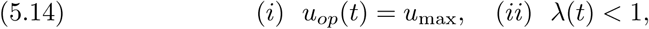

in the interval *t*_*s*_ < *t* ≤ *T* for some *t*_*s*_ < *T*.

Since *R*(*t*) > 0 for all *t* in [0,*T*], we must convert at *u*_max_ at terminal time; otherwise we can always increase *E*(*T*) by converting at a higher rate. The optimal control (5.14) follows from continuity of state and adjoint functions, proving part (*i*).

Upon observing (*i*), the Hamiltonian (5.6) becomes

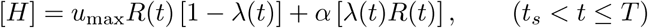

which requires *λ*(*t*) < 1 for *t* in the interval (*t*_*s*_,*T*] adjacent to the terminal time *T* (since the Hamiltonian would be maximized by *u*_max_ in that interval) otherwise. This proves part (ii) of Property 1. (We note parenthetically that the Euler boundary condition (5.8) was not needed for the proof of either part of this Property,)

#### Property 2

If (*t*_*s*_,*T*] is the largest interval adjacent to *T* in which (5.14) holds for *u*_*α*_ > 0 and 0 < *t*_*s*_ < *T*, then *t*_*s*_ must be the zero of *λ*(*t*) = 1 nearest to *T*, giving us the *switch condition*

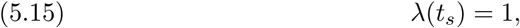

for determining the *switch point t*_*s*_ of the optimal control *u*_*op*_(*t*).

#### Property 3

In the interval (*t*_*s*_,*T*] where the Hamiltonian is maximized by *u*_max_ (with *λ*(*t*) < 1 for *u*_*α*_ > 0), the adjoint DE (5.7) for *λ*(*t*) simplifies to

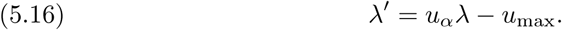

It follows that *λ*(*t*) *is monotone decreasing with time* in (*t*_*s*_,*T*] given *λ*(*t*) < 1 in that interval. The property is also a consequence of the exact solution

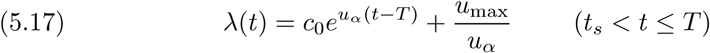

of (5.16). The constant of integration *c*_0_ is determined by the Euler BC (5.8) so that

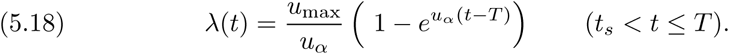

#### Property 4

With *λ*(*t*) increasing as *t* decreases, there exists an instant *t*_*s*_ < *T* when (5.15) holds. The “switch condition” (5.15) determines the switch point *t*_*s*_ to be

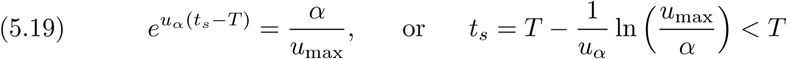

given 0 < *α* < *u*_max_. Hence, the switch point occurs prior to the terminal time (though both are still unknown prior to the application of the threshold condition (5.3)).

We are now in a position to state the following key result on the optimal control:

#### Proposition 5.

*The optimal conversion strategy is u*_*op*_(*t*) = *u*_max_ (0 ≤ *t* ≤ *T*) *if u*_max_ ≤ *α and*

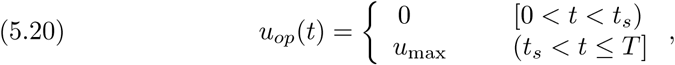

*if u*_max_ > *α, where t*_*s*_ *is determined by (5.19). (Note that t*_*s*_ ↓ 0 *as u*_max_ ↓ *α.)*

*Proof*. The optimal control *u*_*op*_(*t*) must be the lower corner control 0 at least in a small interval (*t*_0_, *t*_*s*_] adjacent to the switch point. If not and *u*_*op*_(*t*) = *u*_max_ for 0 ≤ *t*_0_ < *t* ≤ *t*_*s*_, then

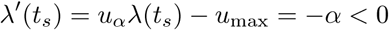

so that *λ*(*t*) is a decreasing function of *t* in some small neighborhood of *t*_*s*_. With 1 − *λ*(*t*) ≤ 0 for *t* ≤ *t*_*s*_, the upper corner control *u*_max_ does not maximize the Hamiltonian at least for *t* in that neighborhood; hence, *u*(*t*) = *u*_max_ is not optimal there. Since the singular solution does not apply, we are left with the only option of *u*_*op*_(*t*) = 0 in that neighborhood. In that case, the adjoint DE (5.7) and the continuity of the adjoint function require

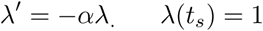

and therewith

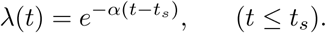

With this, the Hamiltonian,

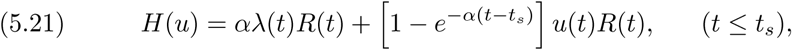

is maximized by the lower corner control *u*_*op*_(*t*) = 0 for all *t* in the interval [0, *t*_*s*_). □

#### Remark 2.

*When u*_max_ ≫ *α, it would seem advantageous not to convert immediately near the start and allow the RB population to grow initially for a while. The larger RB population at a later time would then be converted at the fastest pace possible leading to a larger terminal EB population. Proposition 5 is a realization of this advantage*.

### 5.6. A Conversion Holiday at the Start

That the optimal conversion strategy from our constrained optimization model with a time varying control turns out to be a bang-bang control (5.20) for *α* < *u*_max_ is rather remarkable given the experimental findings in [3]. The relevant data (averaged over several inclusions) reported in there are summarized in Table I.

**Table I:** Mean Count of RB and EB (averaged over inclusions)

| *Bodies* \ *hpi* | 12 | 16 | 20 | 24 | 28 | 32 | 36 | 40 | > 40 |
| --- | --- | --- | --- | --- | --- | --- | --- | --- | --- |
| *RB* | 1.3 | 7.6 | 34 | 105 | 385 | 507 | 271 | 171 | not reported |
| *EB* | 0 | 0 | 0 | 3.7 | 192 | 656 | 706 | 751 | not reported |

It is seen from this summary that only RB chlamydiae (in the two different forms) are observed in the infected cell 20 *hours post infection* (hpi). The pic-charts in Figure 1 of [3] show only RB (including the Dividing RB denoted by DB) particles in the infected cells 20 hours post infection (hpi). With data collected every 4 hpi, the conversion to EB begins sometime before or around 24 hpi. This is well after the initial infection of the host cell, over a time period nearly half of the average duration (over many inclusions) between initial infection and the host cell lysing ending the development cycle.

The total EB population (the sum of the two forms of EB actually observed) rises quickly to 192 units at 28 hpi giving a conversion rate that much greater than the natural RB proliferation rate with every indication of a discontinuous conversion rate with a finite jump discontinuity at the switch point *t*_*s*_ ≲ 24 hpi. Furthermore, with RB population declining sharply after reaching a peak of 507 units at 32 hpi, the corresponding gain in EB units becomes slower as shown by the available data up to 40 hpi. These observations provide further support for conversion being proportional to the RB population size as modeled by (5.5) and at a rate considerably higher than the natural growth rate for the RB form.

It would be natural to embark at this point on the task of parameter estimation (for *α, u*_max_, etc.) and then the determination of the switch point *t*_*s*_ and terminal time *T*. The available data from [3] suggest that our second model still needs further refinement before doing so. From the growth dynamics of the RB population modeled by (5.2), we see for the *α* < *u*_max_ case, that the RB population grows exponentially prior to the onset of conversion (since *u*_*op*_(*t*) = 0 during the conversion holiday at the start) but begins to decline immediately with the onset of conversion since *R*′ = *α* − *u*_max_ < 0 (for *t* > *t*_*s*_). On the other hand, the total RB population (the sum of the two different forms of RB) reported in [3] continues to increase for a period after the appearance of EB particles around 24 hpi. That total only starts to decline after reaching a maximum of 507 units at 32 psi. To remedy this discrepancy, we consider in the next section a refinement of the present second model that allows for two different forms of both RB (denoting the second form by DB) and EB (denoting the pre-EB, intermediate phase by IB) as actually observed in [3]. For linear growth and conversion dynamics with time-varying conversion rate similar to the second model, the optimal conversion strategy for this two-step time-varying conversion model is easily seen to be also bang-bang with a conversion holiday at the start. But unlike the second model, the total RB+DB population in this new third model will be shown to grow for a period of time after the onset of conversion for a realistic range of parameter values and thereby enable us to remove the discrepancy of the second model on a lack of growth after the onset of conversion.

Before leaving the present second model, we note one positive feature among the results for this model that should persist upon further refinements of the second model. It can be that among consequences of the bang-bang solution (5.20) for the model with time-varying control are the following expression for the switch point *t*_*s*_ with

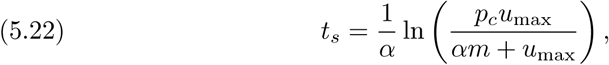

and the terminal time *T* with

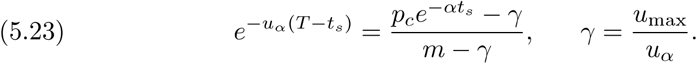

The relation (5.22) shows the switch point is always positive (since we have typically *p*_*c*_ = *P*_*c*_/*N* ≫ *m* + 1 > 1 + *αm*/*u*_max_) and a monotone increasing function of *p*_*c*_ with

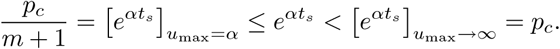

Upon using (5.22) to eliminate *t*_*s*_ from (5.23), we obtain the optimal terminal time

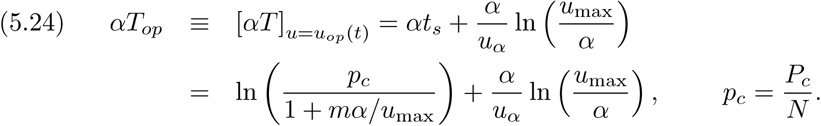

With *u*_max_ > *u*_*α*_ > *α*, the second term is *O*(1) at most, the positive time *T*_*op*_ is then of the order

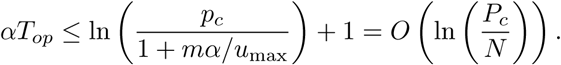

The time to the optimal expected terminal EB population is an order of magnitude smaller than *αT* = *O*(*P*_*c*_) for *u*(*t*) = *α*.

## 6. A TWO-STEP GROWTH AND CONVERSION MODEL

### 6.1. A Realistic Multi-Form Population Model

The separation of Chlamydia population into two distinct populations, reticulate bodies and elementary bodies, were made in the two previous models to enable us to arrive at a relatively simple mathematical model for the purpose of illustrating the mathematics involved in the quantitative approach to one type of issues concerning Chlamydia life cycle. In reality, both reticulate and elementary bodies exist in more than one form. In addition to the RB and EB, two sets of intermediate bodies in transition stage have been observed as the RB convert to EB [3].

A more realistic model for C. trachomatis differentiation and proliferation would be as shown in the schematic diagram in Figure 2:

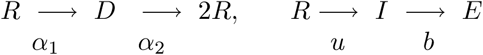

**Figure 2:** A schematic diagram of a two-step duplication and conversion process.

It models the actual observation that the proliferation of RB units is in two phases. At any instant in time, some RB units of its population *R*(*t*) are converted to an *Intermediate form I*(*t*) at the rate *uR* (that would eventually become EB) while the remainder of *R*(*t*) are transformed into a new form, designated as *Dividing RB* at the rate *α*_1_*R*(*t*). Each unit of the population *D*(*t*) of the Dividing RB is capable of binary division into two new RB at the rate *α*_2_*D*. The population *I*(*t*) of Intermediate form converts to EB at a rate *bI*. The concurrent growth rates are summarized mathematically by the following four differential equations:

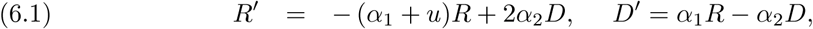

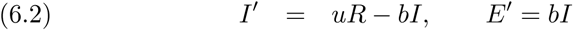

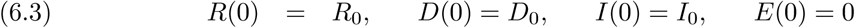

with Dividing RB and Intermediate form populations typically taken to be zero initially.

From the second ODE in (6.2), we get the total EB population at the time of host cell death *T*

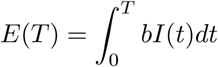

The optimal conversion rate from RB to EB to result in the largest possible EB population at the time of the host cell death time may be stated as the following optimal control problem:

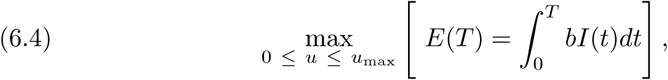

subject to the growth dynamics of the first three ODE and the first three initial conditions of (6.1)- (6.3) and the inequality constraints (5.4). In the context of the approach of the previous sections, some condition for determining the terminal time *T* would also be prescribed.

### 6.2. The RB Population after the Onset of Conversion

Given that all population change rates are linear, the optimal control is expected to be bang-bang for *u*_max_ greater than the natural proliferation rate. Prior to the onset of conversion at *t*_*s*_, we have *u*_*op*_(*t*) = 0 with the first two ODE in (6.1) summed to give

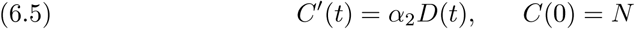

for *C*(*t*) = *R*(*t*) + *D*(*t*) and *R*(0) + *D* (0) = *R*_0_ + *D*_0_ = *N* (with *R*_0_ and *D*_0_ typically taken to be *N* and 0, respectively). With *D*(*t*) > 0 for *t* > 0, we have *C*′(*t*) > 0 so that the total RB population increases with time at least up to the onset of the RB-to-EB conversion.

For *t* > *t*_*s*_, we have from the bang-bang control *u*_*op*_(*t*) = *u*_max_ so that

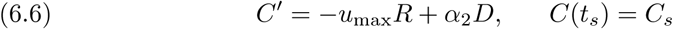

with *C*_*s*_ = *R*_*s*_ + *D*_*s*_ where *R*_*s*_ and *D*_*s*_ are, by continuity, the values of *R* and *D* at the switch point determined by the IVP defined by (6.1) with *u* = 0 and the first two initial conditions in (6.3). Evidently, we have also *C*′(*t*) > 0 (and the total RB population continues to increase) for a period after the onset of the RB-to-EB conversion (as shown by the data collected in [3]) if − *u*_max_*R*_*s*_ + *α*_2_*D*_*s*_ > 0. This confirms the possibility of a two-step model capable of rectifying the remaining deficiency of the highly idealized one step model of the previous section.

The rather crude estimate of the capability of further RB+DB growth in the two-step model after the start of conversion can be further refined:

#### Proposition 6.

*The combined RB+DB population C*(*t*) *increases for t in some interval* (*t*_*s*_, *t*\*) *if α*_2_*D*_*s*_ > *u*_max_*R*_*s*_.

*Proof*. Rewrite the ODE in (6.6) as

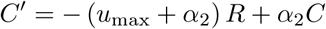

to obtain the unique solution

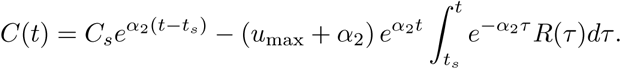

for the IVP. Given the hypothesis, we have by continuity *α*_2_*D* > *u*_max_*R* for all *t* in some interval (*t*_*s*_, *t*\*) adjacent to *t*_*s*_. Within that interval, we have

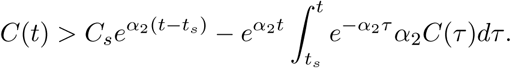

If *C*(*t*) should decrease in that interval, we would have

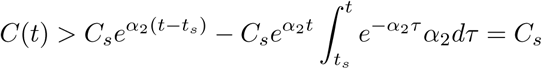

contradicting the assumption and the proposition follows □

As a consequence of the proposition above, the combined RB+DB population *C*(*t*) should continue to increase (at least for an interval of time post-conversion) if the constitution of the bacteria should have *α*_2_*D*_*s*_ > *u*_max_*R*_*s*_ at the switch point when the RB-to-EB conversion begins. Hence, a glaring qualitative discrepancy between the highly idealized model of the previous section can be rectified by the multi-phase life cycle model of this section. The actual (rather complex) details of the corresponding life cycle will be reported in a future work. Instead, we formulate and analyze in the next section a different type of stochastic model to investigate another stochastic aspects of the data reported in [3].

## 7. UNCERTAIN HOST DEATH

### 7.1. The Expected Terminal *EB* Population

Empirical data collected in [3] show that the host cell lysis time does vary for different membrane inclusions, with no lysing reported up to 40+ hpi [3] and lysing observed as late as 72 hpi [2]. A different but realistic approach to investigate the spread of C. trachomatis would be to allow for the uncertainty in the lysis time by treating the terminal time *T* as a random variable with a prescribed probability density function (pdf) postulated or estimated from the data available. For the purpose of analysis, we take the initial time *t* = 0 to be the instant after the initial infecting RB population begins to divide and proliferate. Since host cells are known not to lyse before some *T*_1_ > 0, we should let

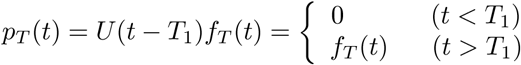

with

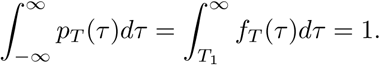

where *U*(*z*) is the Heaviside unit step function with a unit jump at *z* = 0.

The corresponding EB population at terminal time is also a random variable *E*(*T*), a transformed random variable of the random variable *T*. The *expected value* of *E*(*T*), denoted by *E*_*T*_, is given in terms of the pdf *p*_*T*_ (*τ*) by

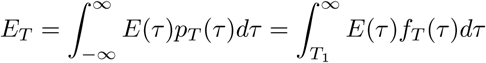

with

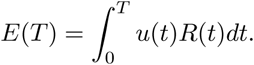

from the growth dynamics (5.2) of EB. For maximum spread of the C. trachomatis bacteria, we choose the (per unit RB particle) conversion rate *u*(*t*) to maximize the expected terminal *EB* population *E*_*T*_:

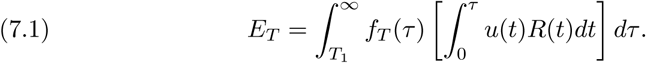

subject to the conversion capacity inequality constraint (5.4). Upon interchanging the order of integration, we may re-write the expression above as

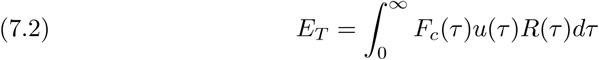

where

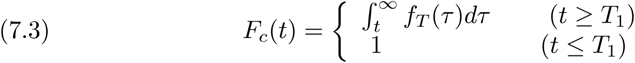

is the probability of the host cell NOT lysed at time *t*. Typical probability density functions include the *uniform density distribution (over the interval* (*T*_1_, *T*_2_))

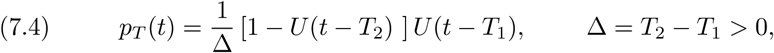

and the *generalized inverse distribution density function*

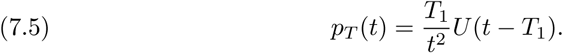

The two corresponding *F*_*c*_(*t*) are

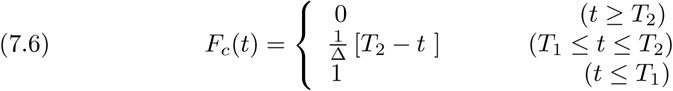

and

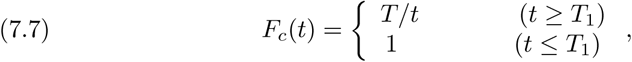

respectively. Given the empirical data available, the uniform pdf with compact support (such as (7.4) leading to (7.6)) would seem a more reasonable characterization of the probability density function for *T*. We will work generally with a pdf with compact support henceforth so that

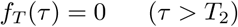

and

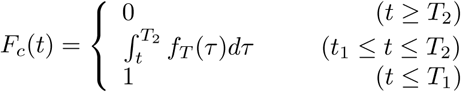

The optimization of expected terminal EB population then takes the form

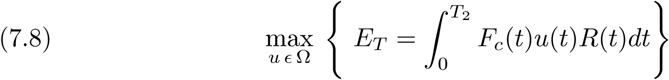

subject to the IVP (5.2) and the constraints (5.4) with Ω being the set of admissible controls as previously defined.

To apply the method of optimal control, we introduce the Hamiltonian

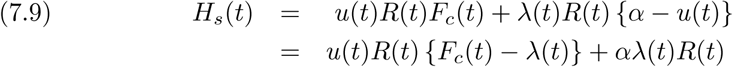

with the new Hamiltonian *H*_*s*_ in (7.9) reduced to the Hamiltonian *H* for a known *T* if *f*_*T*_ (*t*) = *δ*(*t* − *T*) where *δ*(*x*) is the Dirac delta function. The *adjoint function* (aka *Lagrange multiplier*) *λ*(*t*) is to satisfy the adjoint ODE

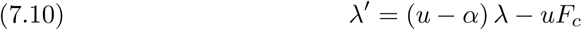

for *t* in the bounded time interval (0, *T*_2_) with an adjoint (Euler) boundary condition (BC) at *T*_2_

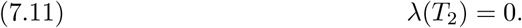

The Maximum Principle requires that we choose an admissible *u* to maximize *H*_*s*_ [1, 4]:

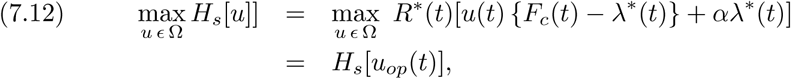

subject to the inequality constraints

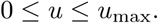

with 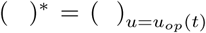. Since the Hamiltonian is linear in the control function *u*(*t*), the optimal control is expected to be a combination of *singular controls* and *extreme values* of the admissible control over different time intervals.

### 7.2. Singular Solution Not Applicable

Candidates for the maximizers of *H*_*s*_ are among the stationary points of the Hamiltonian, i.e., the solutions of

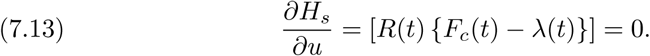

With *R*(*t*) changing at the rate (*α*− *u*)*R* ≥ (*α*− *u*_max_)*R*, the RB population remains positive for all time as long as *N* > 0 so that the stationary condition requires

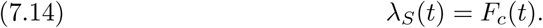

Note that the control *u*(*t*) does not appear in the stationary condition (7.13) and hence is not determined by (7.13). For that reason the immediate consequence of the stationary condition (7.14) is the *singular solution λ*_*S*_(*t*) with the corresponding singular control *u*_*S*_(*t*) to be deduced from other requirements of the Maximum Principle when appropriate. For our problem, the adjoint DE requires

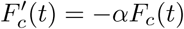

which is generally not satisfied by the probability distribution *F*_*c*_(*t*) of the host cell not lysing (and certainly not by the two illustrative examples in (7.6) and (7.7)). Hence, the singular solution is generally not applicable in any interval of the solution domain [0, *T*_2_] and we have the following negative result for our problem:

#### Proposition 7

*The singular solution (7.14) is not applicable in any part of the solution domain* [0, *T*_2_] *for our uncertain terminal time problem*

It follows from Proposition 7 that the optimal control can only be a combination of the two corner controls 0 and *u*_max_.

### 7.3. Conversion at Maximum Rate for *u*_max_ ≤ *α*

For the range *u*_max_ ≤ *α* or, equivalently,

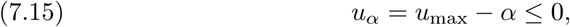

we have the following rather obvious result:

#### Proposition 8

*For u*_*α*_ ≤ 0, *u*_*op*_(*t*) *must be u*_max_ *for all t in* [0, *T*_2_] *with the optimal expected terminal EB population given by*

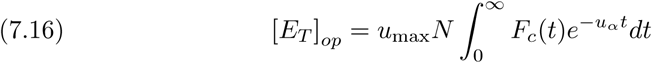

*Proof*. While a more formal proof can be given, it suffices to note that, with *α* − *u*_max_ ≥ 0, the RB population does not decrease even when the conversion to EB is at the maximum rate possible. Converting at any lower rate than the maximum rate would result in less terminal EB and more terminal RB that could have been converted with the highest conversion rate *u*_max_*R* any time in the interval [0, *T*_2_]. □

For the uniform pdf (7.4) with the corresponding *F*_*c*_(*t*) given in (7.6), it is straightforward to calculate from (7.8) the corresponding optimal terminal expected EB:

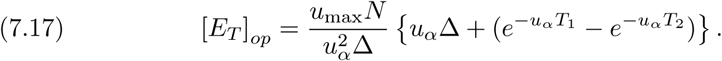

Having dealt with the range *u*_max_ ≤ *α*, we can now focus our discussion of this problem on the complementary and more interesting range *u*_max_ > *α*.

### 7.4. Upper Corner Control Adjacent to Terminal Time

With *R*(*t*) > 0, *u*(*T*_2_) < *u*_max_ is not optimal since we can always choose a larger *u* (still within the admissible range) to convert some of the remaining RB for a larger EB population at *T*_2_. With *u*_*op*_(*T*_2_) = *u*_max_, we have from the adjoint DE

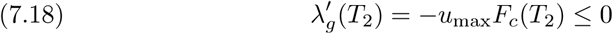

where *λ*_*g*_(*t*) denotes the adjoint function with *u*(*t*) = *u*_max_. Along with the continuity of *λ*(*t*) − *F*_*c*_(*t*), the condition (7.18) requires that *u*_*op*_(*t*) must be *u*_max_ for some interval (*t*_*s*_, *T*_2_] adjacent to *T*_2_. We state this observation more formally in the following proposition with a formal proof:

#### Proposition 9

*The upper corner control maximizes the Hamiltonian (7.9) for some interval* (*t*_*s*_, *T*_2_] *with t*_*s*_ *being the largest root of the switch condition (nearest to T*_2_)

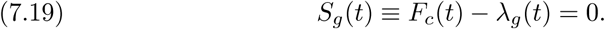

*Proof*. With the Euler BC *λ*_*g*_(*T*_2_) = 0, the Hamiltonian reduces to

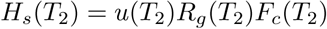

i. For the general case with *F*_*c*_(*T*_2_) > 0, *H*_*s*_(*T*_2_) is maximized by the upper corner control so that *u*_*op*_(*T*_2_) = *u*_max_. Given

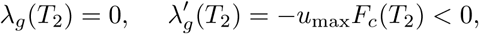

we have *λ*_*g*_(*t*) ≥ 0 but, by continuity, *λ*_*g*_(*t*) < *F*_*c*_(*t*) for some interval (*t*_*s*_, *T*_2_] adjacent to *T*_2_. It follows from *R*(*t*) > 0 that *u*_*op*_(*t*) = *u*_max_ at least in (*t*_*s*_, *T*_2_] with *t*_*s*_ being the root of (7.19) nearest to (but still <) *T*_2_.
ii. For the case *F*_*c*_(*T*_2_) = 0 (and *F*_*c*_(*t*) > 0 for *t*< *T*_2_), we have

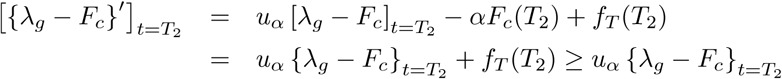

so that *λ*_*g*_ − *F*_*c*_ is an increasing function of *t* but remains < 0 at least in some interval (*t*_*s*_, *T*_2_]. It follows that we have *u*_*op*_(*t*) = *u*_max_ at least in (*t*_*s*_, *T*_2_] with *t*_*s*_ being the root of (7.19) nearest to (but still <) *T*_2_. □

Given Propositions 8 and 9, it remains to determine for the *u*_*α*_ = *u*_max_ − *α* > 0 range: *i*) the optimal control in the complementary range [0, *t*_*s*_), *ii*) the switch point *t*_*s*_; and *iii*) the optimal expected terminal EB population *E*_*T*_. For these tasks, we need the adjoint function for the entire solution interval [0, *T*_2_]. We illustrate the method of the solution for these problems by working out in the next section the details for the uniform density function (7.4).

## 8. UNIFORM DENSITY ON A FINITE INTERVAL

### 8.1. The Adjoint Function

For the uniform probability density function (7.4) and the corresponding *F*_*c*_(*t*) distribution (7.6) for a cell not lysed at time *t*, the adjoint function for the upper corner control *u*_max_ adjacent to the terminal time, denoted by *λ*_*g*_(*t*), is determined by the terminal value problem:

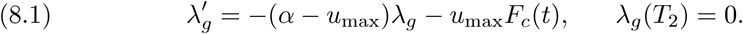

The solution for a finite *T*_2_ and any *t* in the interval [0, *T*_2_] is

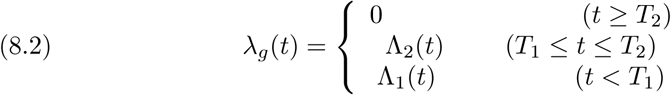

where

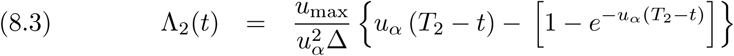

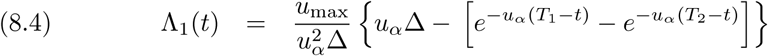

with Δ = *T*_2_ − *T*_1_. Note that *λ*_*g*_(*t*) is continuous at *T*_1_ and *T*_2_. That *λ*_*g*_(*t*) = 0 for *t* ≥ *T*_2_ reflects the fact that the reticulate bodies have no “(shadow) value” beyond *T*_2_. With *F*_*c*_(*t*) = 0 for *t* ≥ *T*_2_, the host cell has already lysed with probability 1 so that we would only be interested in the range of time *t* < *T*_2_.

### 8.2. Lower Corner Control for *t* < *t*_*s*_

In view of Proposition 8, we only need to focus on the range *u*_*α*_ > 0. In that case, we know from Proposition 9 that the upper corner control *u*_max_ is optimal for (*t*_*s*_, *T*_2_) where *t*_*s*_ is the largest zero of (7.19). If *u*_max_ should also be optimal for *t* ≲ *t*_*s*_ for the uniform pdf (7.4), then the ODE for the adjoint function at *t*_*s*_ (the zero of (7.19)) simplifies to

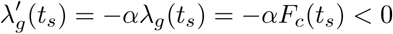

since *λ*_*g*_(*t*_*s*_) = *F*_*c*_(*t*_*s*_). With *λ*_*g*_(*t*_*s*_) a exponentially decreasing function of *t* for *t* ≲ *t*_*s*_ and *F*_*c*_(*t*) a linearly decreasing function of *t*, we have

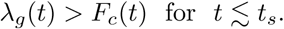

contradicting *u*_max_ being optimal there. Since the singular control is not applicable, this leaves us with the lower corner control as the only option. and we have the following result for the optimal control:

#### Proposition 10

*For u*_*α*_ > 0, *the lower corner control maximizes the Hamiltonian in* [0, *t*_*s*_) *so that the optimal control is the bang-bang control*

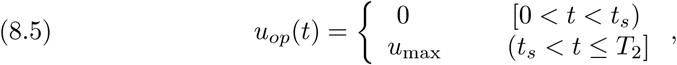

*where t*_*s*_ *is given by switch condition*

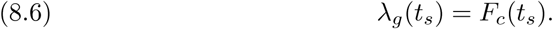

*Proof*. We have already *u*_*op*_(*t*) = *u*_max_ for *t*_*s*_ < *t* ≤ *T*_2_ from Proposition 9 where the *switch point t*_*s*_ is the root of (8.6) nearest to *T*_2_. We also learned from the development prior to this proposition that *u*_*op*_(*t*) cannot be *u*_max_ or the singular solution for *t* ≲ *t*_*s*_.

For the lower corner control in that range, the corresponding adjoint function, denoted by *λ*_*ℓ*_(*t*), is determined by

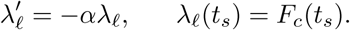

The exact solution is

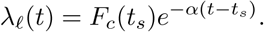

For *t*_*s*_ ≤ *T*_1_, we have *F*_*c*_(*t*) = 1 and 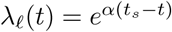 so that 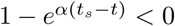 for all *t* < *t*_*s*_ and

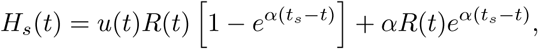

is maximized by *u*_*op*_(*t*) = 0 there.

For *t*_*s*_ > *T*_1_, *F*_*c*_(*t*) decreases only linearly with increasing *t* while *λ*_*ℓ*_(*t*) decay exponentially with *λ*_*ℓ*_(*t*_*s*_) = *F*_*c*_(*t*_*s*_) so that *u*_*op*_(*t*) = 0 is also optimal for *t*< *t*_*s*_. □

### 8.3. The Switch Point *t*_*s*_

The process of determining the switch point *t*_*s*_ for *u*_max_ > *α* depends on the location of *t*_*s*_ since the expressions for *F*_*c*_(*t*_*s*_) and *λ*_*g*_(*t*_*s*_) vary with the range of *t*_*s*_ (see (8.2)).

#### 8.3.1. *t*_*s*_ ≤ *T*_1_

The conditions (8.2) and (8.4) requires

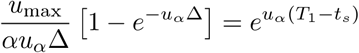

so that

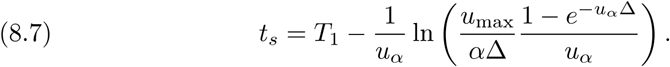

As *u*_max_ increases (with *α* fixed), the switch point *t*_*s*_ tends to *T*_1_ and may have already moved into the *t*_*s*_ > *T*_1_ range (to be discussed below). At the other extreme, the switch point *t*_*s*_ in this range tends to 0 (or less) as *u*_max_ ↓ *α* from above (as we would expect given maximum conversion for *u*_max_ ≤ *α* by Proposition 8).

#### 8.3.2. *T*_1_ < *t*_*s*_ ≤ *T*_2_

For *t*_*s*_ in the interval (*T*_1_, *T*_2_), we have from (??) and (8.2)

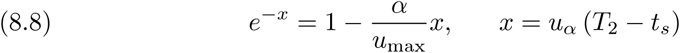

Unlike the case *t*_*s*_ ≤ *T*_1_, the switch condition does not determine *t*_*s*_ explicitly. Instead, it only leads to a nonlinear equation for *t*_*s*_. Graphing the two sides of (8.8) as functions of *x* with *α* and *u*_max_ prescribed shows that the switch condition determines a unique positive root *x*\* with

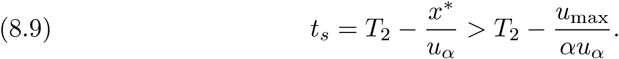

bound above by *T*_2_ and below by *T*_2_ − *u*_max_/ (*αu*_*α*_). For a fixed *α*, we have

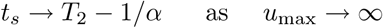

while *t*_*s*_ should move into the *t*_*s*_ < *T*_1_ range as *u*_max_ ↓ *α*.

### 8.4. The Optimal Expected Terminal EB Population

#### 8.4.1. A Local Maximum

For *u*_*α*_ > 0, we know from Proposition 10 that the bang-bang control (8.5) maximizes the Hamiltonian with *t*_*s*_ determined by (8.9) or (8.7) whichever is appropriate. (If *u*_*α*_ ≤ 0, then the optimal strategy for maximizing the expected terminal EB population is to convert at the maximum rate possible from the start as the argument for Proposition 8 applies.) To the extent that maximizing Hamiltonian is not synonymous with maximizing *E*_*T*_, we still need to prove that *u*_*op*_(*t*) maximizes *E*_*T*_ for *u*_*α*_ > 0.

##### Proposition 11.

*For u*_*α*_ > 0, *the bang-bang control (5.20) maximizes the expected terminal EB population*

*Proof*. From Propositions 9 and 10, we know already that *u*_*op*_(*t*) as given by (5.20) maximizes the Hamiltonian *H*_*s*_ with *t*_*s*_ in the interval (*T*_1_, *T*_2_). For our relatively simple control problems, we may appeal to the uniqueness of the switch point and that the optimal control is superior to not-converting any RB to EB at all. □

#### 8.4.2. The Expected Terminal EB Population

For the optimal conversion rate *u*_*op*_(*t*) given in (5.20), we have from the growth dynamics (5.2)

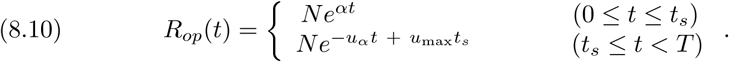

The associated expected EB population is given by (7.8)

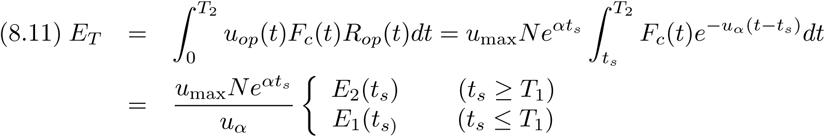

with

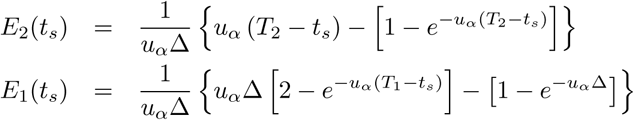

where Δ = *T*_2_ − *T*_1_ and *t*_*s*_ determined by (8.9) if *t*_*s*_ ≥ *T*_1_ or (8.7) if *t*_*s*_ ≤ *T*_1_.

## 9. CONCLUDING REMARKS

We have embarked on a project of modeling and analysis to investigate the proposition that the rather unusual life cycle of C. trachomatis is a consequence of the bacteria’s effort for Darwinian survival. More specifically, we postulate that this effort takes the form of maximizing the EB population at the terminal time of the life cycle with the RB-to-EB conversion rate being the controlling instrument for this effort. To the extent that there appears to be considerable uncertainty in the features of the bacterial developmental cycle, including the terminal time of the development, we formulate and analyze several probabilistic models to determine the optimal RB-to-EB conversion rate that maximizes the (expected) terminal infectious EB population, each with a specification of the terminal time.

The first is a birth and death process type model with time-invariant probabilities for RB population growth and RB-to-EB conversion with a terminal time determined by different mechanisms for termination of development, including a critical weighted sum of the total RB and EB population that reflects the substantial size difference between the RB and EB units. It is found that the expected terminal EB population is maximized by setting the conversion probability equal to the RB growth probability *α*_*D*_. This first model serves as a stepping stone for further refinements toward more appropriate models. Clearly relaxing the time-invariant requirement on the conversion probability *α*_*C*_, the instrument for optimization, could lead to a larger (expected) terminal EB population for faster spread of the infectious disease.

The simple step of permitting the conversion probability to be a function of time *t* transforms the mathematical problem of maximizing the expected terminal EB population to a rather complicated problem in optimal control. For the more interesting case with the maximum allowable conversion rate higher than the natural growth of the RB population, an immediate payoff is an optimal conversion rate that calls for no conversion over a finite time period immediately after the infection of a new host cell. The conversion holiday at the start of a life cycle is precisely what is reported in the empirical findings of [3]. In addition, the duration of the life cycle is now an order of magnitude shorter than the unrealistically long terminal time dictated by the *α*_*C*_ = *α*_*D*_ conversion strategy.

There is however one qualitative deviation of the model prediction from the observed life cycles to be addressed. The RB population from the model declines immediately after the onset of conversion, without the period of continual RB population growth reported in [3]. The discrepancy is then shown to be an artifact of lumping the two different forms of RB units in the idealized model. By a more realistic third model that retains the two known forms of both RB and EB units, we establish the existence of a period of continual growth whenever there is a finite DB (duplicating RB) population at the start of the RB-to-EB conversion.

As the true cause on the life cycle termination is not known and the duration of a life cycle has been observed to vary (see [3]), we undertake an investigation of another kind of models that does not prescribed a specific criterion for determining the terminal time *T* (when the host cell lyses). Instead, we consider *T* to be a random variable with a prescribed probability density function (pdf) taken to be representative of the available data on the host cell lysing time. The stochastic optimal control problem for a pdf uniformly distributed over a finite time interval (*T*_1_, *T*_2_) was worked out in detail. The optimal control for the new problem is again bang-bang with the corresponding RB declining immediately after the onset of RB-to-EB conversion. This discrepancy is to be resolved in a sequel to this paper by a multi-form model similar to the one analyzed in an earlier section herein.

